# Multi-dimensional predictions of psychotic symptoms via machine learning

**DOI:** 10.1101/2020.03.02.974246

**Authors:** Jeremy A Taylor, Kit Melissa Larsen, Marta I Garrido

**Author notes:** **Correspondence** Jeremy Taylor, Melbourne School of Psychological Sciences, University of Melbourne, Redmond Barry Building, Tin Alley, Parkville, Victoria, Australia.

## Abstract

The diagnostic criteria for schizophrenia comprise a diverse range of heterogeneous symptoms. As a result, individuals each present a distinct set of symptoms despite having the same overall diagnosis. Whilst previous machine learning studies have primarily focused on dichotomous patient-control classification, we predict the severity of each individual symptom on a continuum. We applied machine learning regression within a multi-modal fusion framework to fMRI and behavioural data acquired during an auditory oddball task in 80 schizophrenia patients. Brain activity was highly predictive of some, but not all symptoms, namely hallucinations, avolition, anhedonia and attention. Critically, each of these symptoms was associated with specific functional alterations across different brain regions. We also found that modelling symptoms as an ensemble of subscales was more accurate, specific and informative than models which predict compound scores directly. In principle, this approach is transferrable to any psychiatric condition or multi-dimensional diagnosis.

## 1. Introduction

Schizophrenia diagnoses comprise a diverse range of heterogeneous symptoms which collectively manifest in widespread neuroanatomical and functional differences. Over the past decade, machine learning has been widely used in the field of neuroimaging for mapping symptomatic manifestations onto brain substrates. These methods are generally considered a potential gateway to precision psychiatry as they provide predictions at the individual level, hence going beyond classical univariate methods which can only tell us about overall group effects within a given population. The vast majority of psychiatric machine learning studies are primarily concerned with group membership, in particular the binary classification between patients and controls, patient subgroups, or prognoses (1–3). However, given the wide array of symptoms which characterise schizophrenia, individuals each present with their own distinct set of symptoms despite having the same categorical diagnosis. In an effort to parse these symptomatic differences, there has recently been a shift away from dichotomous labels towards a dimensional approach (4, 5).

As defined by the Diagnostic and Statistical Manual of Mental Disorders (DSM; 6), symptoms are categorised as *positive* or *negative*. Positive symptoms are typically absent in the general population, such as hallucinations and delusions, whereas negative symptoms present more often, including affective flattening and poverty of speech. In clinical practice, standardised psychometric tools are widely used to assess the severity of symptoms, in particular the Scale for Assessment of Positive Symptoms (SAPS; 7), the Scale for Assessment of Negative Symptoms (SANS; 8) and the Positive and Negative Symptom Scale (PANSS; 9). Individual symptoms or *subscales* are assigned numeric scores relative to their severity, ranging from absent to severe. The *composite* score is the sum of all symptom subscales, providing an overall summary of the given category. The limited prior work on predicting schizophrenia symptoms via machine learning has thus far only been performed on the basis of composite symptoms (10, 11), general functioning (12, 13) and polygenic risk scores for schizophrenia (14). Other neuroimaging studies have also reported univariate correlates (15, 16), or lack thereof (17, 18), with symptom severity on the basis of composite summary scores rather than those of the underlying symptoms, an approach which significantly comprises aetiological specificity (19). For example, if we were to compare two patients, one with disorganised thought processes which render them unable to bathe themselves to another with severe alogia who is unable to communicate, these are vastly different symptoms which in turn are likely to be caused by different sources of dysfunction in different neural networks. By combining the breadth of symptoms under the hypernyms of schizophrenia, positive or negative symptoms, the superposition of features pertaining to each specific symptom may appear more heterogeneous en masse than if these symptoms were addressed separately. Furthermore, a pair of individuals may be assigned the same composite score, and yet have vastly different symptomatology (e.g., a large number of mild symptoms or a smaller subset of high severity symptoms). Given the pervasive issues associated with the heterogeneity of schizophrenia, we suggest the distinction between individual symptoms may be pertinent.

The aim of this study was to predict the severity of schizophrenia symptoms on a continuum using a dimensional diagnosis approach, based on individual neural and behavioural responses to an auditory oddball task performed during a fMRI scan. We applied machine learning regression techniques within a multi-modal fusion framework to predict each individual symptom whilst determining the set of neural and behavioural features which inform each model. In addition, we sought to predict global symptom severity as an ensemble of these subscales and compare this to models which predict the composite scores directly. Finally, we provide maps of the brain regions which contributed toward predictions of specific symptoms.

## 2. Methods

### 2.1 Dataset

The data used in this study was provided by the Mental Illness and Neuroscience Discovery Institute Clinical Imaging Consortium (MCIC; 20) via the Collaborative Informatics Neuroimaging Suite (COINS; 21) online data repository. Anonymised medication data was also obtained directly from the curators of the dataset. These data were originally collected across multiple sites — the University of New Mexico, Massachusetts General Hospital and University of Iowa.

### 2.2 Participants and cognitive characterisation

From an initial sample of 118 schizophrenia patients obtained from the COINS database, participants were excluded on the basis of missing data and/or poor task performance (mean — 3SD). The final sample consisted of 80 schizophrenia patients (58 male, 22 female) with ages ranging from 18 to 60 years (mean ± SD, 32.55 ± 11.39 years). Diagnoses were confirmed using the Structured Clinical Interview from DSM-IV or Comprehensive Assessment of Symptoms and History (22) with severity of symptoms assessed using the SAPS and SANS. For a summary of participant symptom scores, refer to Supplemental Figure S1. Both SAPS and SANS use a five point scale for each subscale (0 = absent, 1 = questionable, 2 = mild, 3 = moderate, 4 = marked, 5 = severe) with the summary score the sum of all subscales.

### 2.3 Stimulus paradigm

The auditory oddball task used in this study examines the interplay between the involuntary orientation of bottom-up attention toward salient novel stimuli and sustaining voluntary top-down attention towards target stimuli (23). fMRI data were acquired whilst participants listened to streams of predictable standard tones (1kHz, *p* = 0.82), interspersed with infrequent target (1.2kHz, *p* = 0.09) and novel tones (complex, computer-generated tones, *p* = 0.09). Participants were instructed to respond to target stimuli via button press whilst ignoring standard and novel tones. All stimuli had a duration of 200ms, presented at inter-stimulus intervals ranging from 550 to 2050ms (mean 1200ms). The total experiment consisted of four blocks with 120 stimuli per block, resulting in a total 480 trials (396 standard, 42 target and 42 novel). The presentation of stimuli and recording of associated reaction times was performed using E-Prime (version 1.1; Pittsburg, Pennsylvania).

### 2.4 MRI data acquisition

Images were acquired at one site using a 1.5T Siemens Sonata MRI scanner, whilst the remaining two sites both used 3T Siemens Trio. All sites shared closely matched acquisition sequences (repetition time, 2s; echo time, 30ms for 3T, 40ms for 1.5T; flip angle, 90°; field-of-view 22cm; in-plane resolution 3.4mm), with resulting volumes (27 slices, 4mm thickness, 1mm inter-slice gap) providing whole brain coverage. For more information on the standardisation, calibration and quality assurance procedures used to minimise variability across sites, refer to Gollub et al. (20).

### 2.5 MRI pre-processing

Image analysis was performed using SPM12 (Wellcome Trust for Neuroimaging, University College London, London) for MATLAB (version 2016a; The MathWorks, Inc., Natick, Massachusetts). Following slice timing correction and spatial realignment, fMRI images were co-registered to skull stripped *T*_1_-weighted images with origins set to the anterior commissure, normalised to MNI space with voxel size 2 × 2 × 2mm, then smoothed using an 8 × 8 × 8mm full width at half-maximum Gaussian kernel. First level analyses were performed by modelling target and novel stimuli as events with standards as an implicit baseline, button presses and movement parameters as nuisance regressors. For each participant, we obtained contrast images for the target and novel conditions.

### 2.6 Features

Four distinct feature sets were defined categorically as neural responses to target and novel conditions, behavioural measures, and other potential confounds.

For the target and novel conditions, voxel-wise activity was extracted from a set of 15 regions-of-interest within the fMRI contrast images. Informed by a meta-analysis by Kim (24), these regions were assumed *a priori* to be those where task-relevant and irrelevant oddball effects would be most robust. The complete set of regions is shown in Figure 1 with atlas references available in Supplemental Table S1.

**FIGURE 1.**
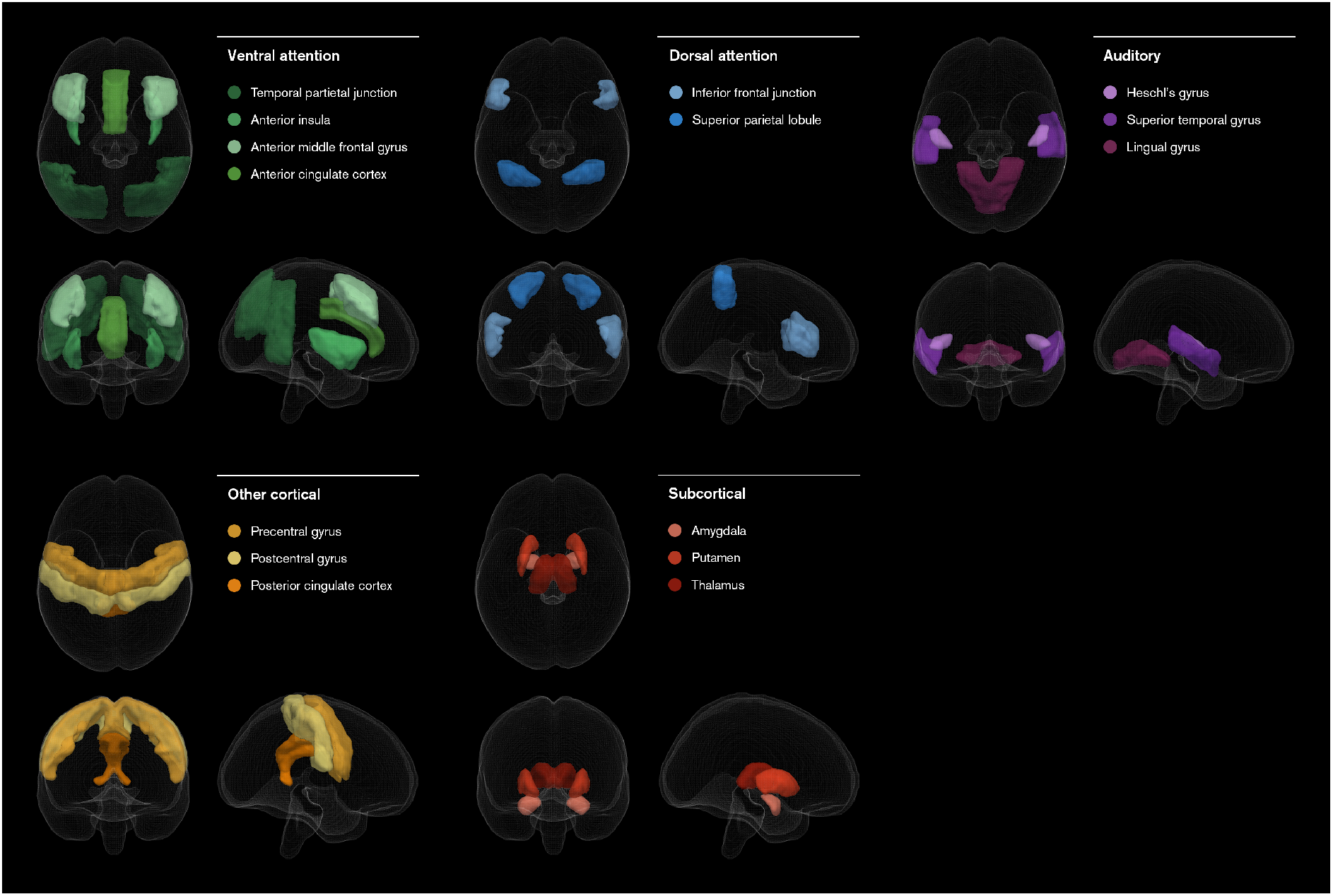
Regions-of-interest and systemic categorisations applied to the fMRI contrasts, defined *a priori* according to a meta-analysis of auditory oddball processing tasks by Kim (24). For Harvard-Oxford cortical and subcortical atlas identifiers, refer to Supplemental Table S1.

The behavioural measures comprised the sensitivity, specificity, precision and mean reaction time of responses to target stimuli, as well as the ex-Gaussian parameters *μ*, *σ* and *τ* (25) summarising the distribution of reaction times over the course of the experiment obtained via the *exgauss* toolbox for MATLAB (version 1.3; 26).

The set of potential confounds comprised age and education, both of which have also been associated with auditory prediction error signals on a univariate basis (17), scanner field strength, which varies between sites, and cumulative antipsychotic drug exposure (20, 27), which is thought to alter neuroanatomy.

### 2.7 Framework

Using these four distinct feature sets as a basis, we employed a machine learning framework which encompasses an *ensemble* of domain *experts* (28). Each expert is trained on a single feature set, which are then integrated using a multi-stage fusion tree as shown in Figure 2.

**FIGURE 2.**
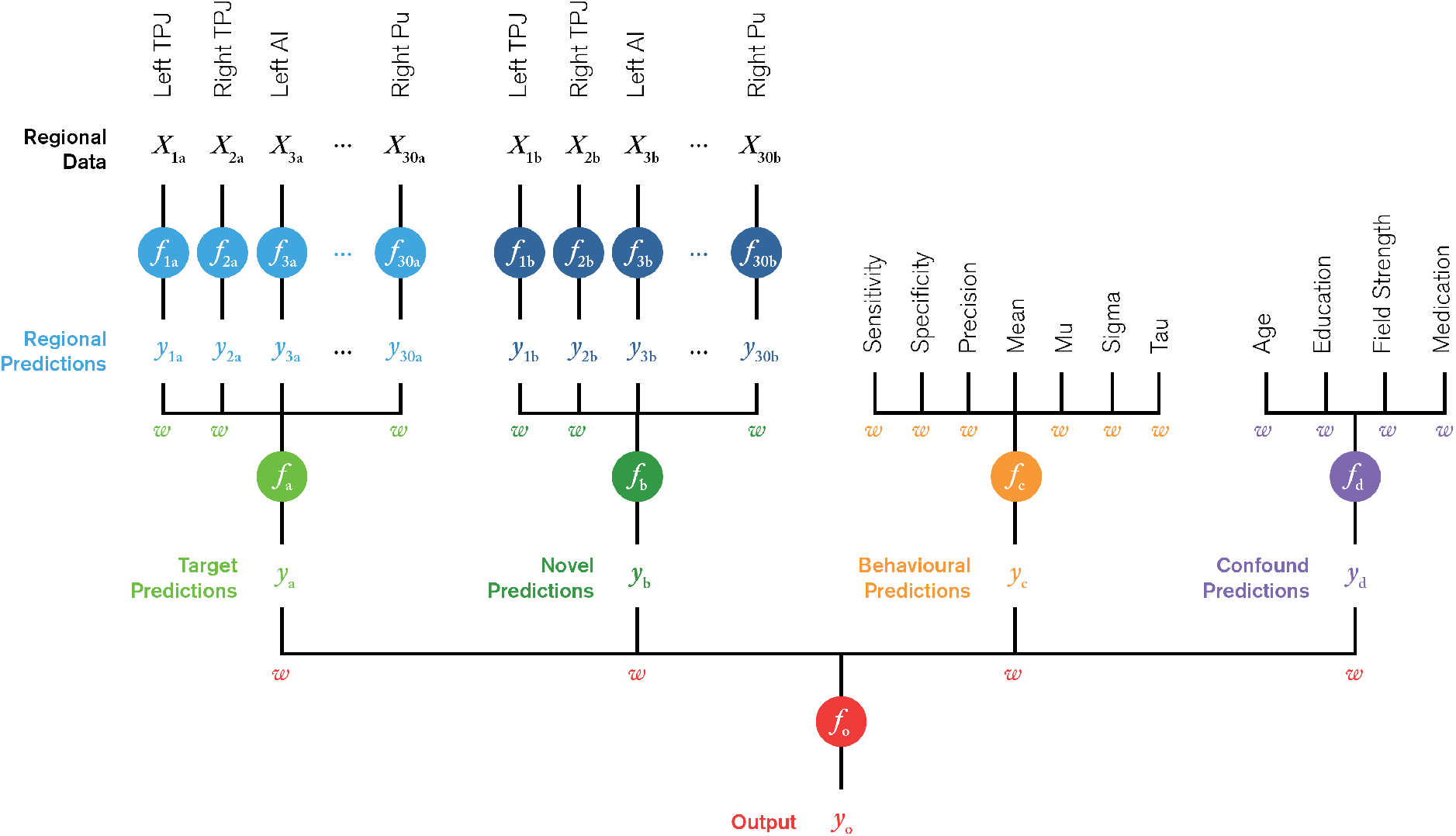
Schematic of machine learning framework. In Stage 1, a set of experts (blue) are independently trained on a subset of fMRI data extracted from one of 30 regions and one of two experimental conditions; target, *a*, and novel, *b*. In Stage 2, the region-based predictions, *y*_1-30a_ and *y*_1-30b_, are then fused to obtain conditional predictions, *y*_a_ and *y*_b_ (green). Experts are also trained on the behavioural (orange) and confound (purple) feature sets. In Stage 3, the conditional, behavioural and confound predictions, *y*_a-d_, are fused to form a final output prediction, *y*_o_ (red). Each fusion model assigns a set of weights, *w*, to each feature set which is used to intuit the relative feature importance in making predictions.

In Stage 1, a set of experts (denoted *f*) were each independently trained on data extracted from a single region-of-interest (ROI), such that each voxel was a feature and each region was a set of features (*X*) with one expert per region. This process was repeated for both target and novel contrasts. Collectively, this set of experts provides a set of region-based predictions (*y*_1-30a_ and *y*_1-30b_) for both the target and novel conditions (subscripts *a* and *b*, respectively).

In Stage 2, these regional predictions were taken as inputs to a pair of secondary fusion models (*f*_a_ and *f*_b_) which return a single conditional prediction (*y*_a_ and *y*_b_) and a set of weights assigned to each region (*w*_1-30a_ and *w*_1-30b_). Experts based on the behavioural and confound feature sets (*f*_c_ and *f*_d_) are also introduced, again assigning weights to each feature.

In Stage 3, the conditional, behavioural and confound predictions (*y*_a-d_) are fused to form a final prediction (*y*_o_) and output weights were assigned to each of the categorical feature sets (*w*_a-d_).

This framework was used to train a set of models to predict each of the symptom subscales and summary scores outlined in Table 1. A late fusion approach was then applied to the subscale predictions by creating summary score ensemble models, taking the predictions from each of the constituent subscale models and fusing them to obtain overall predictions of the SAPS and SANS.

**TABLE 1.** Summary of model performance. Avolition, anhedonia, attention and hallucinations yielded best performance for individual symptoms, whilst the ensemble of subscales outperformed singular summary score models. All *p*-values were computed via 1000 permutations with Bonferroni correction for multiple comparisons.

|  |  |  | *R* | *p*-value |  | *MSE* | *p*-value |  |
| --- | --- | --- | --- | --- | --- | --- | --- | --- |
|  |  |  |  | uncorrected | Bonferroni |  | uncorrected | Bonferroni |
| Individual dimensions | Negative symptoms | Affective Flattening | 0.03 | 0.374 | 1.000 | 0.071 | 0.030 | 0.390 |
|  |  | Alogia | -0.15 | 0.893 | 1.000 | 0.049 | 0.504 | 1.000 |
|  |  | Avolition | 0.59 | 0.001 | 0.013 | 0.049 | 0.001 | 0.013 |
|  |  | Anhedonia | 0.52 | 0.001 | 0.013 | 0.050 | 0.001 | 0.013 |
|  |  | Attention | 0.60 | 0.001 | 0.013 | 0.047 | 0.001 | 0.013 |
|  | Positive symptoms | Delusions | 0.35 | 0.002 | 0.026 | 0.060 | 0.002 | 0.026 |
|  |  | Hallucinations | 0.72 | 0.001 | 0.013 | 0.054 | 0.001 | 0.013 |
|  |  | Bizarre Behaviour | -0.10 | 0.808 | 1.000 | 0.043 | 0.724 | 1.000 |
|  |  | Formal Thought Disorder | 0.40 | 0.002 | 0.026 | 0.046 | 0.001 | 0.013 |
| Summary scores | Single model | SAPS | 0.24 | 0.021 | 0.273 | 0.062 | 0.004 | 0.052 |
|  |  | SANS | 0.39 | 0.001 | 0.013 | 0.019 | 0.001 | 0.013 |
|  | Ensemble model | SAPS | 0.51 | 0.001 | 0.013 | 0.018 | 0.001 | 0.013 |
|  |  | SANS | 0.49 | 0.001 | 0.013 | 0.017 | 0.001 | 0.013 |

### 2.8 Implementation

The machine learning framework, as illustrated in Figure 2, was implemented using the Scikit-learn (version 0.20.2; 29) and NumPy (version 1.15.4) libraries for the Python programming language (version 3.6.5; Python Software Foundation). All experts and fusion models were trained using the *lasso* algorithm (30), which constrains the size of the model coefficients through regularisation and setting a subset of feature coefficients to zero. The result is a sparse formulation with implicit dimensionality reduction and a high level of model interpretability.

Each target score (*y*) was rescaled to a zero to one range, as per the maximum possible score on the given scale. All neuroimaging features were standardised to their respective *z*-scores, rescaling across participants to zero mean and unit variance. Dimensionality reduction was performed using principal components analysis, projecting the data onto a subset of components which explained 90% of the total variance, such that the number of features was much less than the number of samples (*M* ≪ *N*). Non-neuroimaging features were also standardised to the *z*-scores, however, did not require further dimensionality reduction given *M* ≪ *N*.

To make predictions for each individual subject, we employed a 10-fold cross-validation scheme with 10 repetitions, stratified by site with the regularisation hyper-parameter *α* optimised using a nested 9-fold cross-validation scheme. Predictions for the summary score ensemble models were obtained by averaging across those from each of the constituent subscale models.

Model performance was evaluated by comparison of true and predicted scores using the mean-squared error (MSE) and Pearson’s correlation coefficient (*R*). The statistical significance of each model was tested by 1000 permutations of the target variables, with *p* < 0.05 for both metrics indicating that the model has truly learned some pattern within the data, subject to correction for multiple comparisons.

## 3. Results

### 3.1 Predicting individual symptom subscales

These data were found to be predictive of some, but not all symptom subscales, as shown in Table 1. The negative symptoms of avolition, anhedonia and attention all had statistically significant correlations between targets and predictions ranging from 0.52 to 0.60 (*p* < 0.013, Bonferroni corrected), whilst the positive symptom of hallucination had the highest correlation of 0.72 (*p* < 0.013, Bonferroni corrected).

All models had a comparable mean-squared error, ranging from 0.047 to 0.054, which translates to approximately 23% error on the original scale. Although the delusions and formal thought disorder models had seemingly significant correlations of 0.35 and 0.40, inspection of the prediction plots shown in Figure 3 indicates that these models were constrained in their predictions, suggesting a possible bias toward the sample means (0.53 and 0.15, respectively).

**FIGURE 3.**
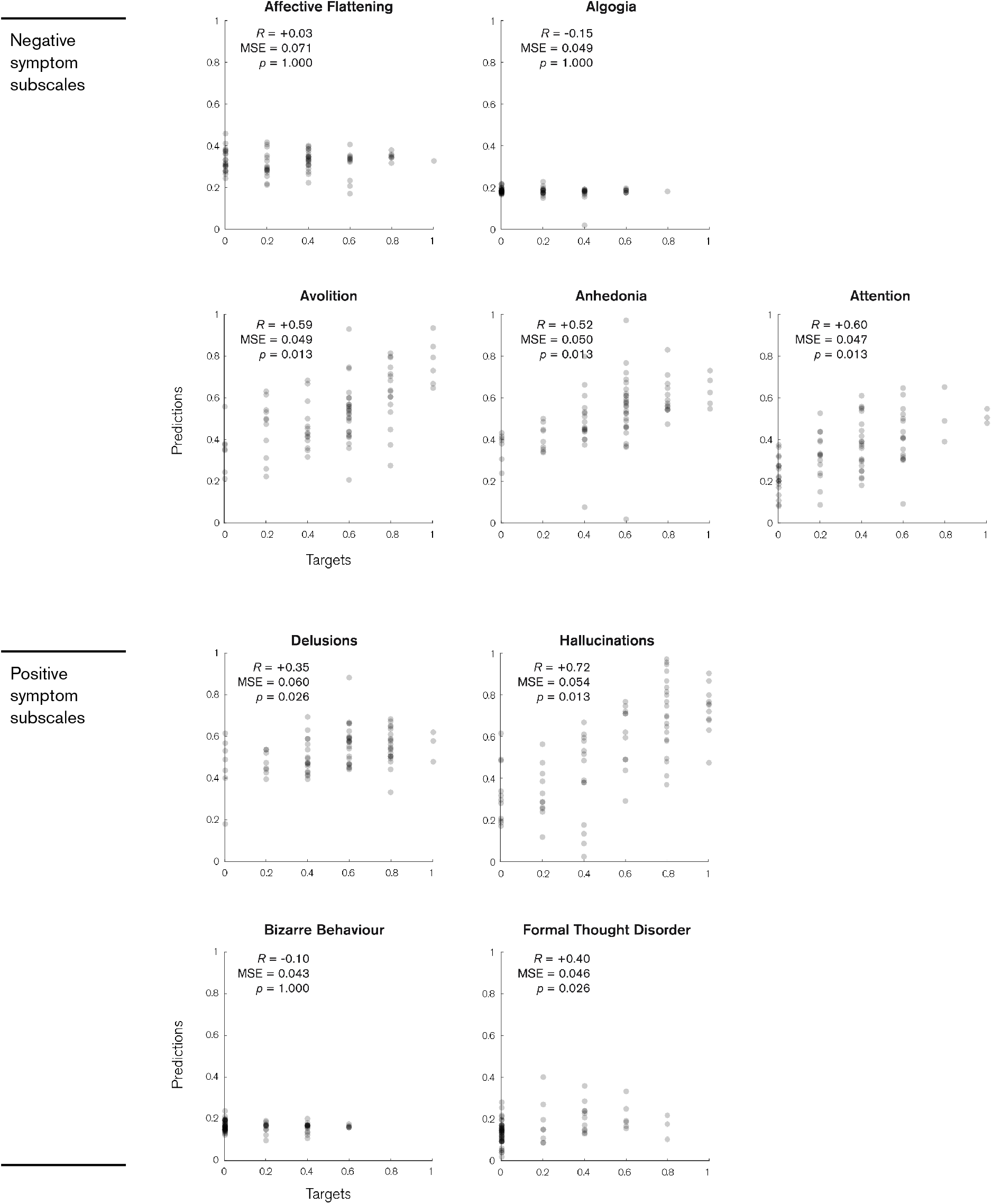
Plots of model predictions and true scores for individual subscales within the SAPS and SANS. Predictions for each subject are shown as grey dots. All scores are rescaled to a zero to one scale. *p*-values are Bonferroni corrected for multiple comparisons.

### 3.2 Predicting symptom summary scores

We applied two different approaches in predicting the SAPS and SANS summary scores. In the first case, a single model was trained directly on the summary scores using the same framework as depicted in Figure 2. As shown in Table 1 and Figure 4, the composite positive symptom model was not statistically significant and the negative symptoms only yield a modest correlation. However, when considering each of the individual subscale models as an expert on a particular symptom within an ensemble which collectively predicts the summary score, we found a marked improvement in performance. Both of these ensemble models proved statistically significant (*p* < 0.013, Bonferroni corrected), with positive symptoms demonstrating a two-fold increase in correlation (0.24 to 0.51) and a decrease in mean-square error (0.062 to 0.018). On average, this translates to an approximate 13.4% error margin.

**FIGURE 4.**
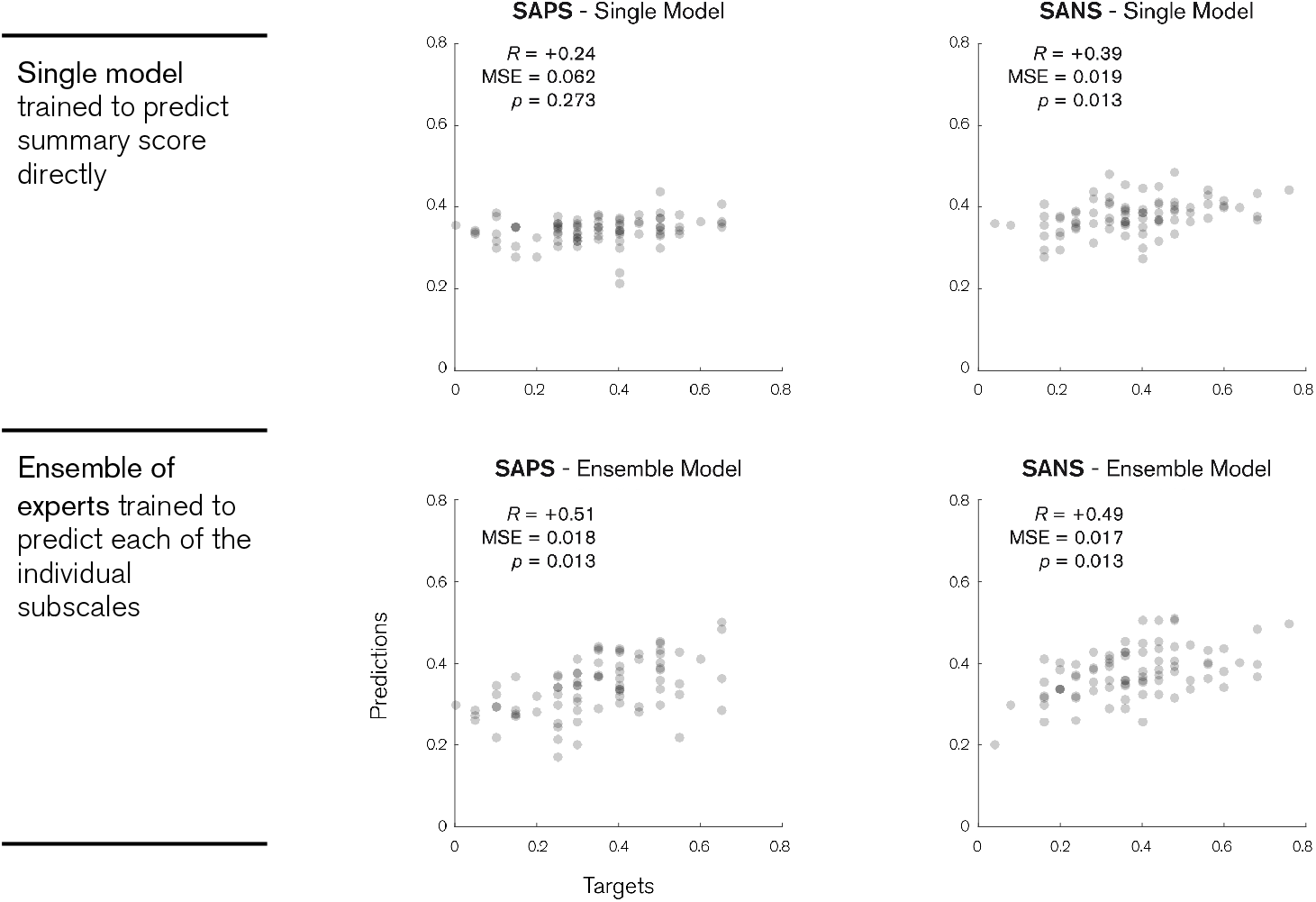
Plots of model predictions and true scores for the SAPS (left) and SANS (right) summary scores. Top row shows the predictions for a single model trained to predict the summary scores directly. Bottom row shows predictions for an ensemble model, where each expert is trained to predict one of the individual subscales. Predictions for each subject are shown as grey dots. All scores are rescaled to a zero to one scale. *p*-values are Bonferroni corrected for multiple comparisons.

### 3.3 Subscale model explanations

Given that we are able to predict a number of symptoms from the available data, we are also able to intuit the main factors which underpin model performance by examining the contributions of each feature set in a *post hoc* manner. This is achieved by examining the fusion weights assigned to each feature set in Stage 3 and individual features in Stage 2.

Firstly, we wish to establish whether neuroimaging is indeed useful, given the added time and monetary investment necessary to acquire these data. The target and novel feature sets were highly weighted in the Stage 3 output fusion (between 72 and 100%) in comparison with the behavioural and confound feature sets (0 to 17% and 0 to 11%, respectively) in each of our statistically significant models. This indicates that the neuroimaging data was the main driver behind the predictions, above and beyond the behavioural data obtained via the task itself. However, excluding those for the anhedonia model, the behavioural and confound coefficients were not set to zero, therefore these features still had some, albeit negligible contribution to the final predictions (Supplemental Table S2).

To better understand the contribution of individual neuroanatomical features within these neuroimaging feature sets, the output weights from Stage 3 were applied to the categorical fusion weights from Stage 2. For each score, we are then able to obtain a regional weight map, as illustrated in the main 4 × 15 matrices of Figure 5a. Here, each element represents a single feature with colour indicating which features were identified by the algorithm as important, and conversely, which were non-informative. In the top panel, each row represents a region-of-interest, with columns indicating the hemisphere (denoted L and R) and experimental stimulus (target and novel). Furthermore, by adding these elements together, we can collapse across subsets of features to summarise the broader system-wide differences between each brain system (4 × 5 matrix), hemisphere (4 × 1 vector) and experimental condition (bottom 2 × 1 vector). For visualisation purposes, Figure 5b shows the regional weight maps projected onto a three-dimensional representation of the original regions-of-interest, collapsed across conditions.

**FIGURE 5.**
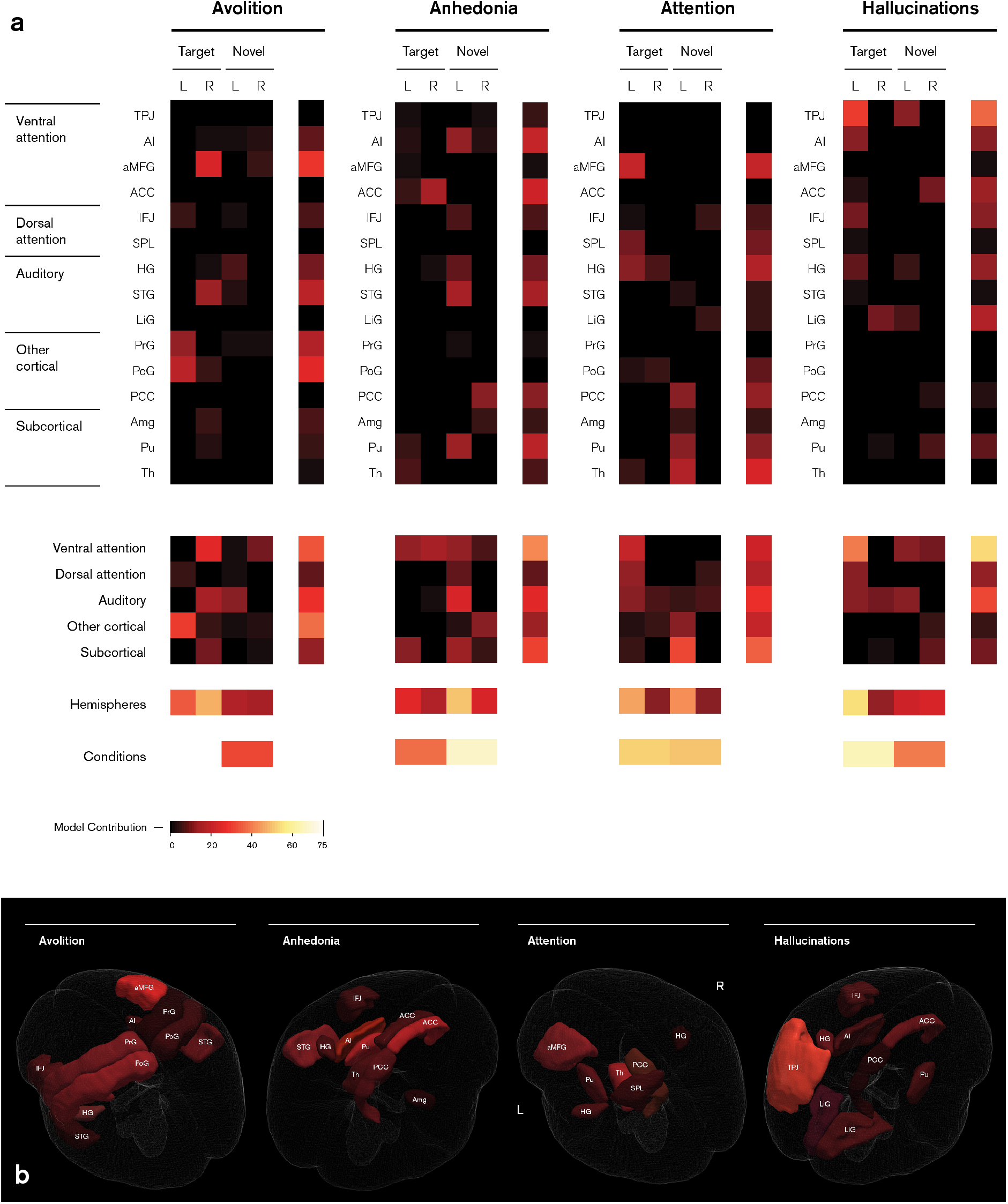
Feature importance for individual subscale models. Colours denote the contribution of each feature toward predictions as a percentage, with black indicating an entire feature set has been marked as irrelevant by the *lasso* algorithm. **(a)** Regional weight maps (top panel) show the relevance of each region (rows) in both left (L) and right (R) hemispheres under target and novel conditions (columns). Categorical weight maps (bottom panel) show the net contribution of each system, hemisphere and condition toward model predictions. Categorical groups of regions are based on Kim (24). **(b)** Regional weight maps projected onto three-dimensional brain structures.

In the avolition model, predictions were mainly driven by the target response, namely the right anterior middle frontal gyrus (aMFG) and other cortical activity in the left hemisphere. For anhedonia, predictions were primarily informed by the anterior cingulate cortex (ACC) target response and left superior temporal gyrus (STG) novel response. Attention predictions were equally driven by both target and novel stimuli, in particular the left aMFG response to target stimuli and subcortical activity in the novel condition. Hallucinations were informed by the left hemispheric target response, principally the ventral attention network and temporal parietal junction (TPJ). Critically, pairwise similarity measures between weight vectors indicated that each symptom had its own distinct pattern of activity across different sets of regions (Supplemental Figure S2).

### 3.4 Summary model explanations

The difference between the two approaches in predicting summary scores is most apparent when comparing the respective weight maps. For the single positive symptoms model (Figure 6), we can observe that of all the possible combinations of features, the optimal solution computed by the algorithm comprises a single feature — the response to target stimuli in the left precentral gyrus. Given the low performance of this model, this can be attributed to a classic case of underfitting. Conversely, the ensemble model weight map includes the specific contributions toward each of the individual subscales, resulting in a widespread distribution of features across the whole brain. Notably, we observe that the precentral gyrus is not a member of the ensemble weight map, nor any of the constituent subscales. Similarly, the negative symptom models demonstrate the same pattern, with the single model reduced to the left inferior frontal junction (IFJ) and right Heschl’s gyrus (HG) responses to the target condition, both of which are down-weighted in the equivalent ensemble model.

**FIGURE 6.**
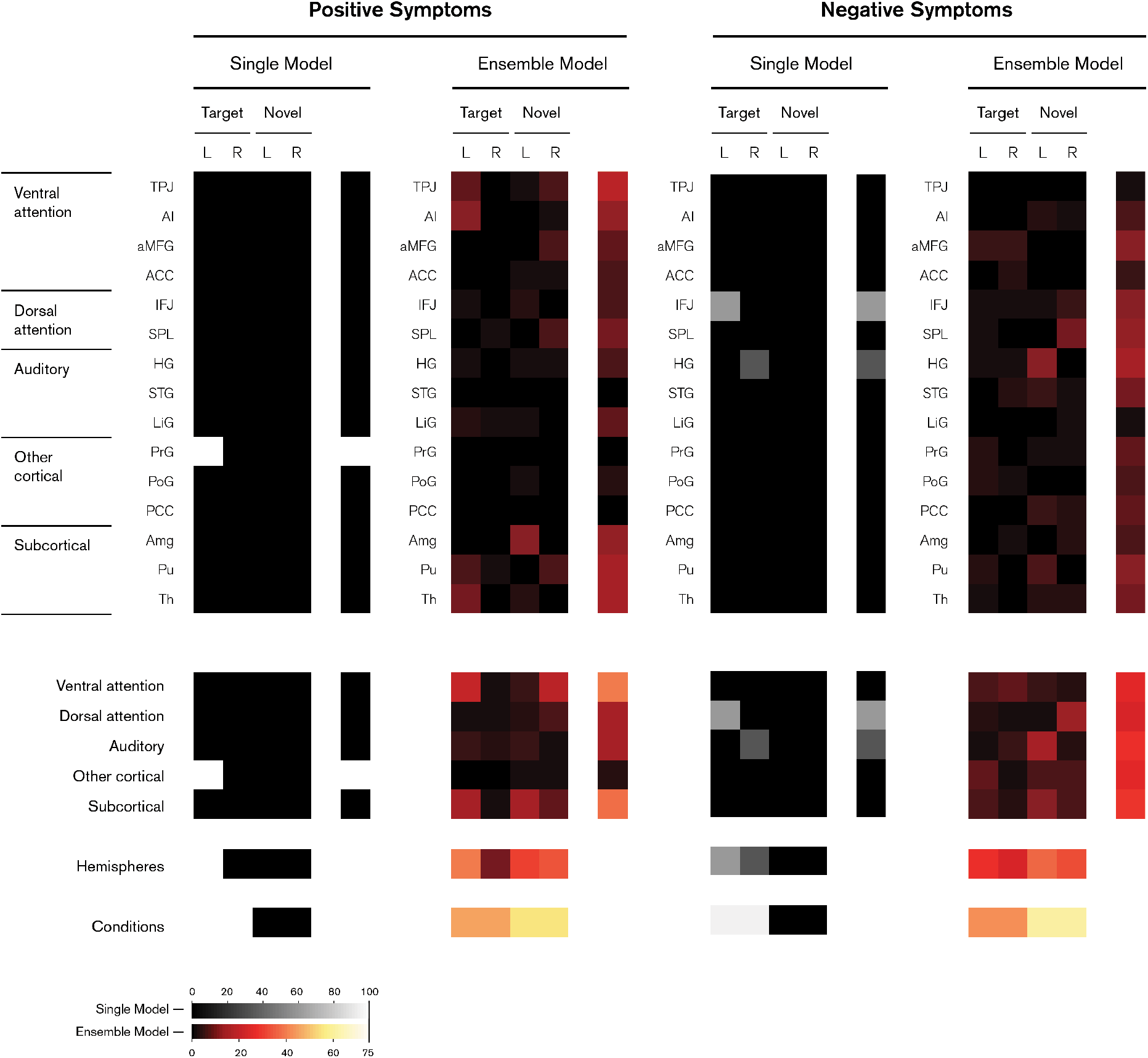
Feature importance for SAPS and SANS summary score models. Ensemble models are shown in warm colour map and single models shown in greyscale. Regional weight maps (top panel) show the relevance of each region (rows) in both left (L) and right (R) hemispheres under target and novel conditions (columns). Categorical weight maps (bottom panel) show the net contribution of each system, hemisphere and condition toward model predictions. Categorical groups of regions are based on Kim (24). Colours denote the contribution of each feature toward predictions as a percentage, with black indicating an entire feature set has been marked as irrelevant by the *lasso* algorithm.

These comparisons demonstrate that the ensemble of subscales clearly outperforms the single model approach, not only in terms of predictive accuracy, but also in terms of identifying plausible functional neuroanatomical maps — the composite negative and positive symptoms arise from widespread brain networks, whereas individual symptoms pertain to more nuanced sub-networks.

## 4. Discussion

In this study, we used multivariate machine learning regression techniques to predict the severity of schizophrenia symptoms on a continuum based on the neural and behavioural responses to an auditory oddball task. By training a set of models to predict each symptom subscale independently, these data were found to be highly predictive of hallucinations, attention, avolition and anhedonia. We also found that by modelling the composite SAPS and SANS summary scores as ensembles of these subscales, the accuracy of predictions significantly increased, whereas single models trained to predict summary scores directly demonstrably underfit to irrelevant features.

### 4.1 Interpreting functional anatomy of psychotic symptoms

Anhedonia is described as a reduced capacity to experience pleasant emotions (31). We found that anhedonia predictions were mainly driven by responses to the novel stimulus in the left hemisphere, in particular the putamen and STG, as well as the target response in the ACC. The ACC is known to play a key role in reward processing (32) and has previously been linked to anticipation of pleasant events (33) and self-referencing (34) in schizophrenia. Deep brain stimulation of the ACC has also been shown to modulate anhedonia-like symptoms (35). To the best of our knowledge, the putamen and STG have not been linked to anhedonia in schizophrenia specifically, however have been previously reported in depression, which has high comorbidity with anhedonia (31). The putamen is connected to the motor cortices and is thought to encode associations between stimuli, actions and rewards (36). Notably, major depressive disorder patients (MDD) with anhedonia have a two-fold age-related putamen volume decrease in comparison with healthy controls (37). Additionally, the STG, primarily involved in auditory and language processing, has a reduced response in first-episode MDD patients with anhedonia when comparing probable vs. improbable rewards (38). STG volume reduction has also been widely reported in schizophrenia patients (39). Collectively, these findings may imply that anhedonia leads to a lower anticipation for rewards upon performing the required action, as reflected in changes within the putamen, STG and ACC.

Predictions of avolition, i.e. a lower pursuit and persistence of goal-directed activities (31), were predominantly informed by the target response in the right aMFG. This region has been previously associated with processing of conflicting information (40) and is reported to have reduced activity under working memory load in those at ultra-high risk for psychosis (41). Target responses in the left precentral and postcentral gyri also contributed to the prediction, albeit to a lesser degree. Alterations in activity within these sensorimotor regions may suggest that those with avolition are required to make an increased effort in response to stimuli which demand a physical action.

Predictions of attention scores were largely informed by the target response in the left aMFG, a region engaged in tasks requiring divided attention (42). In schizophrenia, activity in the left MFG during sustained attention has been previously been shown to correlate with compound negative symptoms on the PANSS scale (43). Patients with brain tumours in the left MFG also show significant reductions in flexible attention and cognition (44). Volitional or self-initiated shifts in attention in the absence of instructional cues have been associated with both left and right MFG activity (45, 46). Interestingly, the thalamic response to the novel condition was also highly weighted, which is known to filter distracting or conflicting information (47, 48). Given that participants were instructed to ignore novel stimuli, the thalamus involvement here may be inhibiting these distractors and allowing for increased selective attention.

Our model for hallucinations was primarily driven by the target response within the ventral attention network and the left TPJ, a known critical node in the speech perception network implicated in hallucinations (49, 50). The left TPJ has previously been used as a target area for transcranial direct-current stimulation (tDCS), leading to a reduction in hallucinations in schizophrenia patients (51). In turn, hallucinations following tDCS have been shown to correlate with the functional connectivity between the left TPJ and left AI (52). This is consistent with our findings, which identify both of these regions as highly predictive of hallucination severity.

Further studies investigating symptom-specific circuitries may open new possibilities for informing future personalised treatments. In the future, if we were to obtain a robust brain mapping for each symptom based on consistent replications of these findings, brain stimulation or pharmacological interventions could be tailored for an individual based on their symptom profile with dosages relative to the level of severity. This could also provide opportunities for reverse translation into an array of symptom-based animal models of schizophrenia.

### 4.2 Methodological considerations

We employed a multi-modal fusion approach which has previously been applied to neuroimaging data for combining different types of structural and functional images (53–55). When building an integrative model from a dataset comprising multiple distinct feature sets, there are two general approaches for fusing these features to form a single prediction. In an *early fusion* approach, features from each modality can be merged prior to the learning process with the joint representation input to a single model providing a multimodal prediction (56). Alternatively, in a *late fusion* approach, a set of models can be trained on each feature set independently and the unimodal predictions from each are combined to form an overall multimodal prediction, typically through averaging or a secondary linear model. The key methodological advance presented in this study is that we perform fusion not only on the basis of the data structure, but also on the *target variables*. The psychometric tools used in clinical practice, such as the SAPS and SANS, are by definition multidimensional — they comprise a set of symptom subscales which are assessed independently, then combined to obtain a composite summary score. Our results suggest that within a machine learning context, a late fusion of subscale predictions as per the original diagnostic framework provides greater specificity and accuracy than the early fusion equivalent of predicting the summary scores directly. Together with the drawn region-based weight maps, this finding also suggests that each symptom has a distinct functional anatomic pattern, whereas models trained to predict positive and negative composite scores directly were unable to capture this nuanced information. In principle, we envisage that modelling the symptom subscales as an ensemble could be applied to any psychiatric disorder.

In a typical neuroimaging context, the number of available features vastly outweighs the number of samples, a phenomenon known as the curse of dimensionality (57). This often leads to an overfitting of model parameters, which in turn may not generalise to new samples (1). To address this issue, we chose to adopt a multi-tier fusion tree which is conceptually similar to a well-established approach known as stacked regression (58). This enabled us to iteratively reduce the dimensionality of the data from the original voxel space whilst also improving model interpretability by expressing the relevant features in more general terms of brain regions and networks. As such, the functional anatomy which informs model predictions is more interpretable by those familiar with the pathology of the disease and relatable to other univariate studies.

Note that whilst some neuroimaging features may have a greater contribution than others, all features with non-zero weights contribute to the model predictions. Although a highly weighted feature is unlikely to be a false positive given adequate signal-to-noise ratio (59), the contribution of any one single feature should be interpreted with caution. For example, although left TPJ activity in the target condition may be of greatest importance in our hallucinations model, this feature alone only contributes approximately 24% toward predictions. Given that the optimal solution for this model also includes 17 other regions, the accuracy would likely decrease without these additional features. By collapsing across conditions, hemispheres or networks, we are able to summarise the contributions of subsets of these features, but cannot fully comprehend the interactions between them within the context of the whole feature set.

### 4.3 Conclusion

In conclusion, we suggest that a shift from dichotomous labels toward a multi-dimensional diagnostic approach which delineates the severity of each individual symptom in a piecewise manner may be more useful for parsing the underlying neurobiological causes. In turn, this framework may help improve our understanding of how specific symptoms relate to functional differences within the brain, in schizophrenia or indeed any other psychiatric disorder.

## Acknowledgements

This work was supported by the Australian Research Council Centre of Excellence for Integrative Brain Function (ARC Centre Grant CE140100007), a University of Queensland fellowship (2016000071) and Foundation Research Excellence Award (2016001844) to MIG. We would like to thank Randy Gollub and Margaret King for providing data, as well as Ilvana Dzafic, Saskia Bollmann and Kelly Garner for discussions on fMRI, and the University of Queensland Research Computing Centre (RCC) for access to high performance computing resources.

