## Supplemental Material for "Multi-dimensional predictions of psychotic symptoms via machine learning"

#### Methods

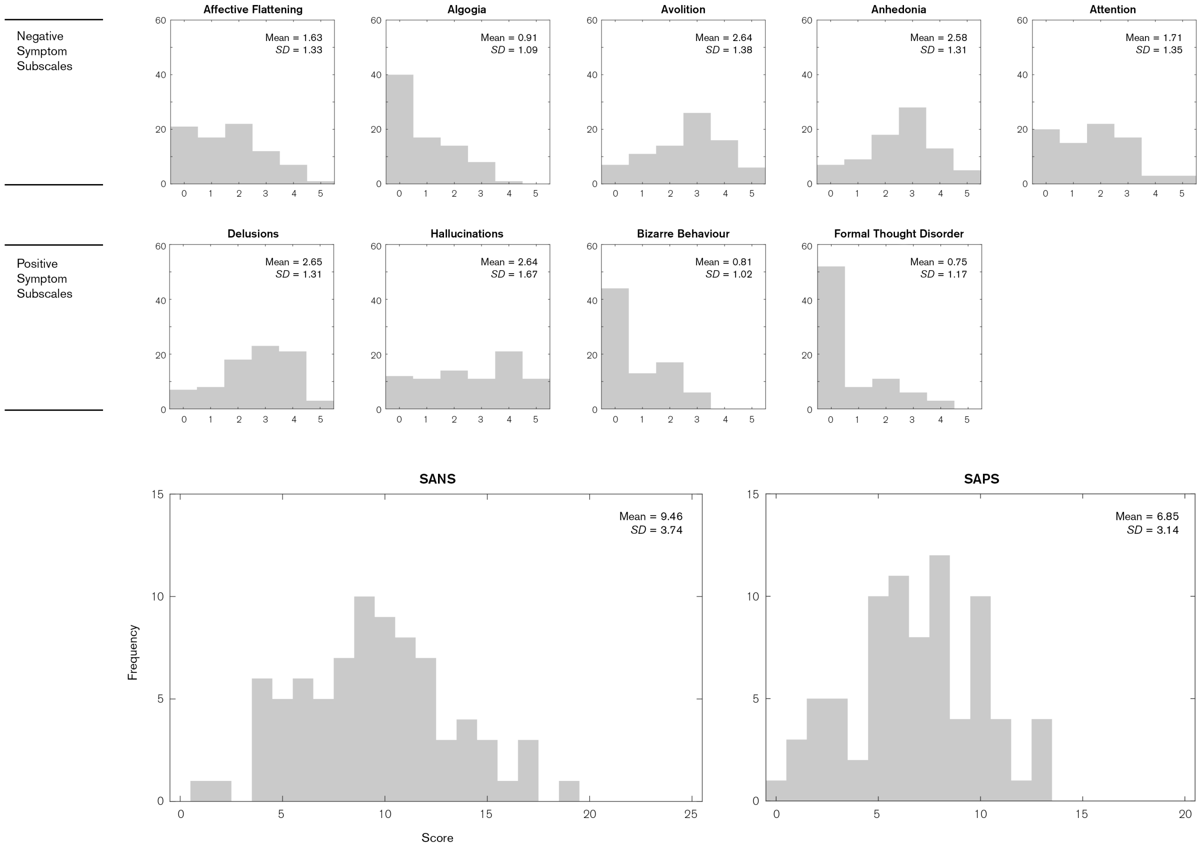


FIGURE S1 — Summary of symptom subscales and summary scores for SAPS and SANS. Each subscale is scored on a 0 to 5 scale with composite scores the sum of all subscales.

TABLE S1 — Regions-of-interest applied to the fMRI contrasts. Anatomical masks for each region were defined using the Harvard-Oxford cortical and subcortical atlases with a probability threshold of 50%. Hemispherical divisions of cortical structures were made using the left and right cerebral cortex as masks. The temporal parietal junction mask was defined as the union of the surrounding regions, as specified by Schurz et al. (1): the lateral occipital cortex, superior division; angular gyrus; supramarginal gyrus, posterior division; middle temporal gyrus, temporooccipital part.

| Classification | Region | Abbreviation | Atlas Label | Atlas ID |
|---|---|---|---|---|
| **Ventral attention** | Temporal parietal junction | TPJ | Lateral occipital cortex, superior division | 22 |
| | | | Angular gyrus | 21 |
| | | | Supramarginal gyrus, posterior division | 20 |
| | | | Middle temporal gyrus, temporooccipital part | 13 |
| | Anterior insula | AI | Insular cortex | 2 |
| | Anterior middle frontal gyrus | aMFG | Middle frontal gyrus | 4 |
| | Anterior cingulate cortex | ACC | Cingulate gyrus, anterior division | 29 |
| **Dorsal attention** | Inferior frontal junction | IFJ | Inferior frontal gyrus, pars triangularis | 5 |
| | | | Inferior frontal gyrus, pars opercularis | 6 |
| | Superior parietal lobule | SPL | Superior parietal lobule | 18 |
| **Auditory** | Transverse gyrus | HG | Heschl's gyrus | 45 |
| | Superior temporal gyrus | STG | Superior temporal gyrus, anterior division | 9 |
| | | | Superior temporal gyrus, posterior division | 10 |
| | Lingual gyrus | LiG | Lingual gyrus | 36 |
| **Other cortical** | Precentral gyrus | PrG | Precentral gyrus | 7 |
| | Postcentral gyrus | PoG | Postcentral gyrus | 17 |
| | Lateral IPS/IPL | | N/A | N/A |
| | Posterior cingulate cortex | PCC | Cingulate gyrus, posterior division | 30 |
| **Subcortical** | Amygdala | Amg | Amygdala | 10/20 |
| | Putamen | Pu | Putamen | 6/17 |
| | Thalamus | Th | Thalamus | 4/15 |
| | Cerebellum | | N/A | N/A |

### Results

TABLE S2 — Output fusion weights for statistically significant subscale models, indicating relative contribution of each feature set at Stage 3 of the framework shown in main manuscript, Figure 1. Values shown are normalised with crosses indicating the entire feature set was marked as irrelevant by the *lasso* algorithm.

| | | Target | Novel | Behaviour | Nuisance |
|---|---|---|---|---|---|
| **Negative symptoms** | Avolition | 0.57 | 0.35 | 0.05 | 0.02 |
| | Anhedonia | 0.42 | 0.58 | × | × |
| | Attention | 0.34 | 0.38 | 0.17 | 0.11 |
| **Positive symptoms** | Hallucinations | 0.49 | 0.34 | × | 0.18 |

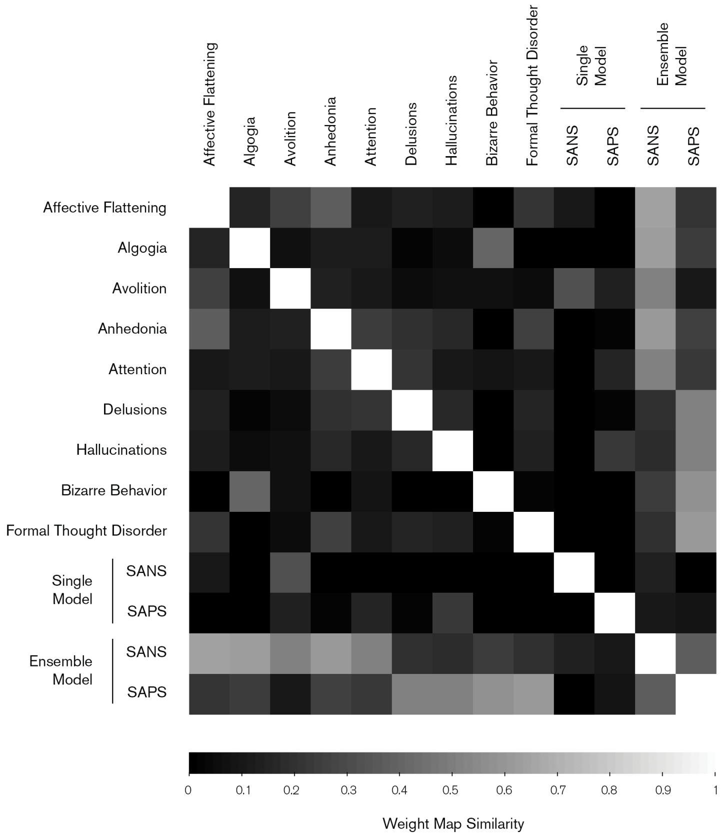


FIGURE S2 — Similarity matrix comparing the weight maps from each model. Pairwise similarities computed as the dot product between weight vectors with values ranging from 0 for dissimilar, orthogonal maps to 1 for similar, collinear maps.
